## Supplementary Material for "Long-read whole genome analysis of human single cells"

### Supplementary Figures

A
B
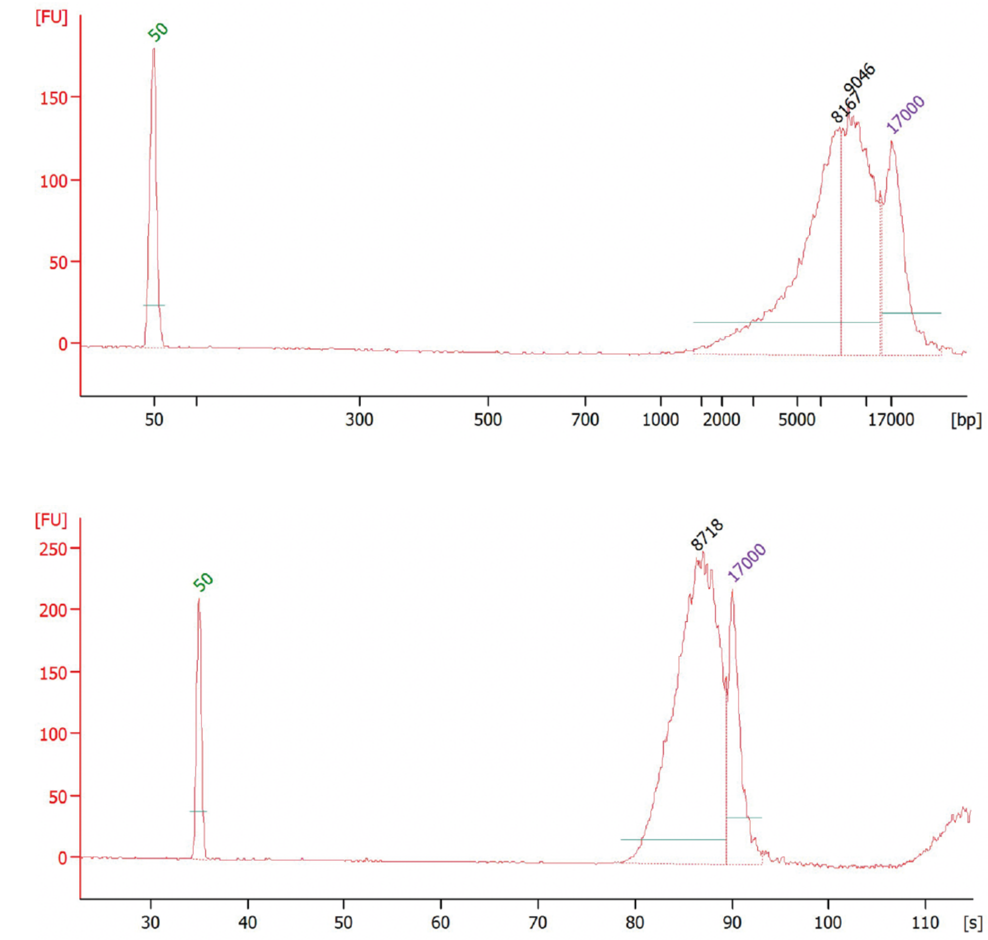

**Figure S1. Fragment length distribution of a single-cell dMDA PacBio library. A)** Bioanalyzer profile of SMRTbell library for T-cell A1 before size selection. **B)** Fragment length distribution of the same SMRTbell library after size selection using AMPure beads.

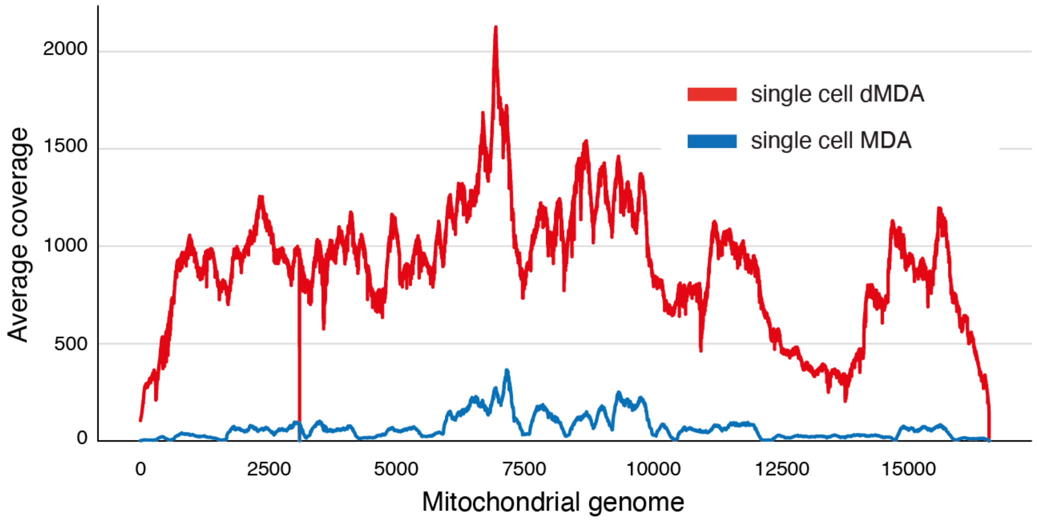

**Figure S2. Average coverage over the mitochondrial genome.** The two lines show the average coverage for the eight Illumina dMDA (red) and eight MDA (blue) single-cell samples.

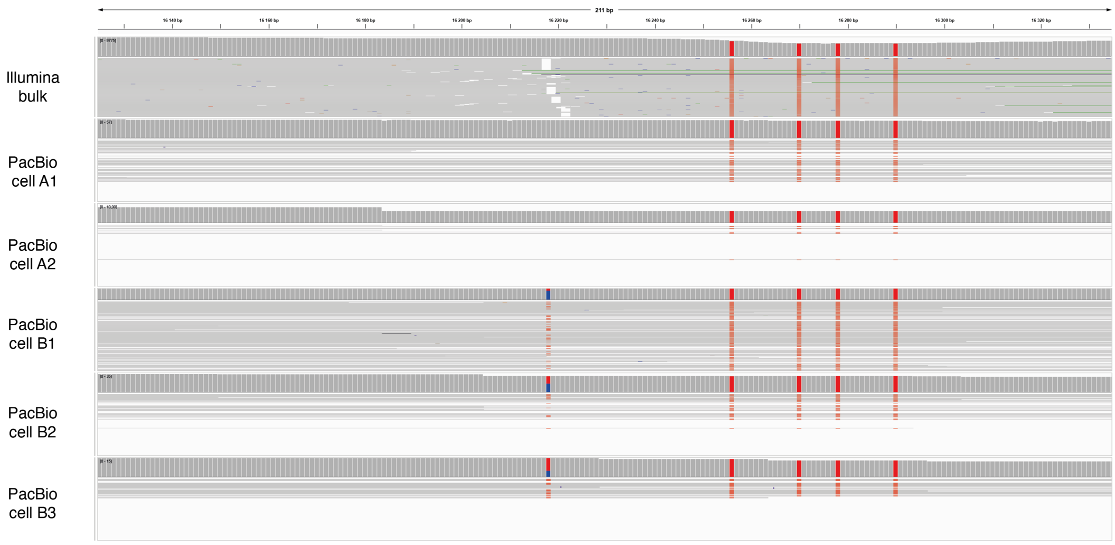

**Figure S3. Mitochondiral heteroplasmy in T-cell B.** IGV plot showing a 211 bp window surrounding position chrM:16,218 where the reference position is a C. In the PacBio data for T-cell B, 41-67% of the reads gave support for a T. In the bulk DNA and in T-cells A1 and A2, 0% of the reads gave support for the T substitution. The four red vertical lines correspond to variants in the mitochondria that are fixed in all three samples.

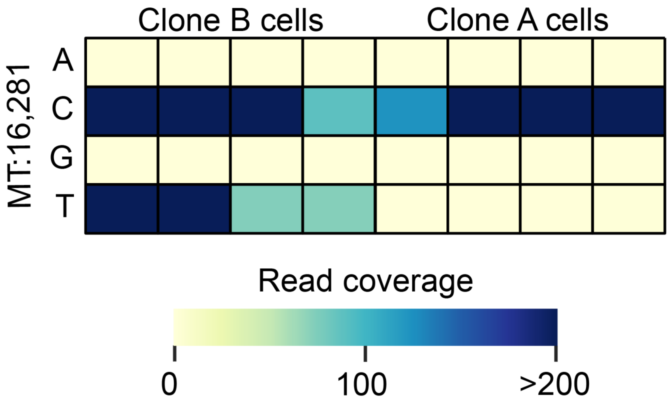

**Figure S4. Validation of mitochondrial heteroplasmy in T-cell clone B.** The heat map shows the distribution of different nucleotides (A, C, G and T) at position 16,281 in T-cell clones B (left) and clone A (right). Only data for the dMDA amplified samples are shown, since several of the MDA samples displayed low or no coverage at this position. All samples for clone B contain both the C and T bases. In the clone A samples only the base C is present.

### Supplementary Tables

**Table S1.** Overview of Illumina WGS data generated in this project.

| ID | Sample | Sample type | Total read pairs |
|---|---|---|---|
| Illumina_dMDA_A1 | T-cell clone A | Single-cell dMDA | 194979790 |
| Illumina_dMDA_A2 | T-cell clone A | Single-cell dMDA | 99366738 |
| Illumina_dMDA_A3 | T-cell clone A | Single-cell dMDA | 118075528 |
| Illumina_dMDA_A4 | T-cell clone A | Single-cell dMDA | 137472548 |
| Illumina_dMDA_B1 | T-cell clone B | Single-cell dMDA | 167428747 |
| Illumina_dMDA_B2 | T-cell clone B | Single-cell dMDA | 163128356 |
| Illumina_dMDA_B3 | T-cell clone B | Single-cell dMDA | 175128648 |
| Illumina_dMDA_B4 | T-cell clone B | Single-cell dMDA | 192217756 |
| Illumina_MDA_A1 | T-cell clone A | Single-cell MDA | 144803454 |
| Illumina_MDA_A2 | T-cell clone A | Single-cell MDA | 171922528 |
| Illumina_MDA_A3 | T-cell clone A | Single-cell MDA | 181849519 |
| Illumina_MDA_A4 | T-cell clone A | Single-cell MDA | 165570616 |
| Illumina_MDA_B1 | T-cell clone B | Single-cell MDA | 123531164 |
| Illumina_MDA_B2 | T-cell clone B | Single-cell MDA | 181450923 |
| Illumina_MDA_B3 | T-cell clone B | Single-cell MDA | 181372742 |
| Illumina_MDA_B4 | T-cell clone B | Single-cell MDA | 198105347 |
| Illumina_Bulk | PMBC Bulk DNA | PCR-Free WGS | 416647211 |

**Table S2.** Statistics for Illumina WGS datasets after random downsampling.

| Sample type | T-cell clone | Downsampled reads | 0x coverage (%)[a] | 5-20x coverage (%)[a] |
|---|---|---|---|---|
| Single-cell dMDA | A | 99976338 | 84.323 | 3.442 |
| Single-cell dMDA | A | 99366738 | 77.627 | 5.627 |
| Single-cell dMDA | A | 99999079 | 78.033 | 5.579 |
| Single-cell dMDA | A | 99994062 | 49.516 | 19.649 |
| Single-cell dMDA | B | 99983362 | 54.081 | 17.350 |
| Single-cell dMDA | B | 99990842 | 70.670 | 9.294 |
| Single-cell dMDA | B | 99996499 | 50.865 | 19.282 |
| Single-cell dMDA | B | 99987004 | 64.442 | 11.810 |
| Single-cell MDA | A | 99986588 | 66.993 | 6.780 |
| Single-cell MDA | A | 99992151 | 81.254 | 5.273 |
| Single-cell MDA | A | 100000818 | 83.715 | 4.369 |
| Single-cell MDA | A | 99985140 | 81.663 | 5.157 |
| Single-cell MDA | B | 100000534 | 72.341 | 4.941 |
| Single-cell MDA | B | 99998491 | 62.896 | 10.812 |
| Single-cell MDA | B | 99987332 | 68.818 | 10.040 |
| Single-cell MDA | B | 99980732 | 94.981 | 1.250 |
| PCR-Free WGS | Bulk | 100002110 | 9.243 | 89.212 |

[a] Percent of bases in the GRCh38 reference that has 0x or 5-20x coverage.

**Table S3.** Overview of PacBio WGS data generated for the five single cells and for the bulk DNA sample.

| Semple name | Sample | Sample type | Total reads | Data amount |
|---|---|---|---|---|
| PacBio single-cell A1 | T-cell clone A | Single-cell dMDA | 2,750,802 | 19.9 Gb |
| PacBio single-cell A2 | T-cell clone A | Single-cell dMDA | 1,561,444 | 15.3 Gb |
| PacBio single-cell B1 | T-cell clone B | Single-cell dMDA | 2,547,184 | 20.2 Gb |
| PacBio single-cell B2 | T-cell clone B | Single-cell dMDA | 1,040,050 | 10.0 Gb |
| PacBio single-cell B3 | T-cell clone B | Single-cell dMDA | 1,306,998 | 13.3 Gb |
| PacBio bulk | PMBC Bulk DNA | HiFi WGS | 5,509,497 | 97.3 Gb |

**Table S4.** PacBio Sequel II run statistics and alignment results for five human single T-cells obtained from the clones A and B

| | Clone A | | Clone B | | |
|---|---|---|---|---|---|
| | **Single-cell A1** | **Single-cell A2** | **Single-cell B1** | **Single-cell B2** | **Single-cell B3** |
| ≥ Q20 reads | 2,750,802 | 1,561,444 | 2,547,184 | 1,040,050 | 1,306,998 |
| ≥ Q20 yield (bp) | 19,880,131,345 | 15,332,739,249 | 20,169,954,798 | 10,010,164,105 | 13,331,286,206 |
| ≥ Q20 read length (mean, bp) | 7,227 | 9,819 | 7,918 | 9,624 | 10,199 |
| ≥ Q20 read quality (median) | Q36 | Q35 | Q36 | Q33 | Q35 |
| Number aligned reads | 2,739,035 (99.57%) | 1,557,468 (99.74%) | 2,517,588 (98.83%) | 1,036,841 (99.69%) | 1,302,581 (99.66%) |
| Number of alignments | 6,508,237 | 5,346,280 | 6,235,924 | 2,713,013 | 3,711,537 |
| Aligned read mean concordance | 99.18% | 99.10% | 99.11% | 99.13% | 98.91% |
| Aligned read length (mean) | 2,994 | 2,785 | 3,149 | 3,620 | 3,525 |
| Aligned read length N50 | 5,429 | 4,821 | 6,476 | 7,391 | 6,539 |
| Aligned read length 95% | 8,691 | 8,867 | 9,739 | 11,636 | 11,387 |
| Aligned read length Max | 43,386 | 41,038 | 48,731 | 51,021 | 54,566 |
| Mean coverage | 6 | 5 | 6 | 3 | 4 |
| Covered bases | 39.60% | 26.95% | 27.71% | 32.98% | 28.86% |

**Table S5.** Overview of SNVs detected in Illumina and PacBio single-cell WGS data

| Sample type | T-cell | Total SNVs | Not found in bulk DNA | Found in bulk DNA[a] |
|---|---|---|---|---|
| Illumina dMDA | A | 407683 | 199999 | 207684 |
| Illumina dMDA | A | 838786 | 224249 | 614537 |
| Illumina dMDA | A | 604022 | 217081 | 386941 |
| Illumina dMDA | A | 2098281 | 344340 | 1753941 |
| Illumina dMDA | B | 1927828 | 322682 | 1605146 |
| Illumina dMDA | B | 1166063 | 249078 | 916985 |
| Illumina dMDA | B | 2132715 | 359900 | 1772815 |
| Illumina dMDA | B | 1562806 | 347843 | 1214963 |
| Illumina MDA | A | 1688193 | 328200 | 1359993 |
| Illumina MDA | A | 886840 | 227187 | 659653 |
| Illumina MDA | A | 791362 | 208210 | 583152 |
| Illumina MDA | A | 850374 | 234589 | 615785 |
| Illumina MDA | B | 1469848 | 278451 | 1191397 |
| Illumina MDA | B | 1895984 | 341829 | 1554155 |
| Illumina MDA | B | 1567015 | 345818 | 1221197 |
| Illumina MDA | B | 336174 | 117117 | 219057 |
| PacBio dMDA | A (A1) | 1934332 | 707971 | 1226361 |
| PacBio dMDA | A (A2) | 1521548 | 737975 | 783573 |
| PacBio dMDA | B (B1) | 1336370 | 474617 | 861753 |
| PacBio dMDA | B (B2) | 1388632 | 432899 | 955733 |
| PacBio dMDA | B (B3) | 1429733 | 525126 | 904607 |

[a] For Illumina data, the comparison was done to Illumina bulk, for PacBio to PacBio HiFi bulk

**Table S6.** Somatic SNVs detected in the two PacBio single-cells for clone A, but not in the PacBio bulk sample or clone B single cells.

| | **Chromosome** | **Position** | **Reference allele** | **Alternative allele** |
|---|---|---|---|---|
| 1 | chr1 | 22628223 | G | A |
| 2 | chr11 | 121519544 | C | G |
| 3 | chr2 | 180266451 | C | A |
| 4 | chr4 | 117567808 | C | G |
| 5 | chr5 | 437214 | G | A |
| 6 | chr8 | 36368389 | C | T |
| 7 | chr8 | 48619816 | C | T |

**Table S7.** Somatic SNVs detected in at least two PacBio single-cells for clone B, but not in the PacBio bulk sample or clone A single cells.

| | **Chromosome** | **Position** | **Reference allele** | **Alternative allele** |
|---|---|---|---|---|
| 1 | chr1 | 113193054 | A | G |
| 2 | chr10 | 34415480 | T | G |
| 3 | chr11 | 75164286 | G | A |
| 4 | chr11 | 78106030 | G | A |
| 5 | chr11 | 105534490 | C | T |
| 6 | chr12 | 124661965 | G | C |
| 7 | chr13 | 85887230 | T | C |
| 8 | chr15 | 74233006 | G | T |
| 9 | chr16 | 71091898 | A | G |
| 10 | chr18 | 37520501 | G | T |
| 11 | chr2 | 48617691 | C | T |
| 12 | chr3 | 151588052 | A | G |
| 13 | chr4 | 474440 | C | T |
| 14 | chr4 | 170294389 | T | G |
| 15 | chr5 | 98938587 | T | G |
| 16 | chr6 | 165180217 | A | T |
| 17 | chr7 | 154618000 | C | T |
| 18 | chr8 | 23441489 | C | T |
| 19 | chr8 | 76709056 | G | A |
| 20 | chr9 | 108593173 | G | C |

**Table S8. Overview of structural variants detected in HiFi data from single cells and bulk DNA.** Only SVs having at least 95% reciprocal overlap with an SV in the bulk DNA sample are reported.

| Sample | INS | DEL | DUP | INV | OTHER[a] | **TOTAL** |
|---|---|---|---|---|---|---|
| pt_022_001(A) | 1015 | 736 | 3 | 6 | 37 | **1797** |
| pt_073_001(A) | 612 | 438 | 1 | 3 | 23 | **1077** |
| pt_027_001(B) | 840 | 597 | 7 | 19 | 37 | **1500** |
| pt_073_002(B) | 421 | 321 | 1 | - | 3 | **746** |
| pt_084_002(B) | 767 | 493 | - | 2 | 7 | **1269** |
| pt_115_001(Bulk) | 13814 | 9679 | 122 | 83 | 503 | **24201** |

[a] DEL/INV, INVDUP and other complex SVs

**Table S9. SVs found in Illumina data.**

| Prep | Clone | Deletions | Tandem_Dup | Inversions | Insertions | BND | Total |
|---|---|---|---|---|---|---|---|
| dMDA | A | 0 | 0 | 0 | 0 | 4 | **4** |
| dMDA | A | 33 | 5 | 1 | 15 | 25 | **70** |
| dMDA | A | 21 | 2 | 0 | 9 | 14 | **44** |
| dMDA | A | 352 | 26 | 5 | 144 | 97 | **584** |
| dMDA | B | 360 | 20 | 3 | 124 | 117 | **580** |
| dMDA | B | 170 | 10 | 3 | 63 | 59 | **283** |
| dMDA | B | 412 | 25 | 6 | 152 | 128 | **674** |
| dMDA | B | 228 | 17 | 2 | 73 | 29 | **373** |
| MDA | A | 19 | 3 | 0 | 11 | 21 | **46** |
| MDA | A | 9 | 1 | 3 | 3 | 10 | **21** |
| MDA | A | 21 | 0 | 1 | 11 | 12 | **40** |
| MDA | A | 14 | 1 | 0 | 8 | 14 | **35** |
| MDA | B | 3 | 1 | 0 | 2 | 4 | **10** |
| MDA | B | 63 | 7 | 0 | 19 | 42 | **123** |
| MDA | B | 56 | 2 | 0 | 17 | 21 | **91** |
| MDA | B | 0 | 0 | 0 | 1 | 4 | **5** |

**Table S10. Number of tandem repeat elements in the single cells having same repeat size as was found in the bulk sample**

| Sample | T-cell clone | Number TRs having same size as in bulk |
|---|---|---|
| PacBio A1 | A | 6512 |
| PacBio A2 | A | 3512 |
| PacBio B1 | B | 4916 |
| PacBio B2 | B | 4504 |
| PacBio B3 | B | 4404 |

**Table S11:** ***De novo*** **assembly results for the two PacBio T-cells A1 and B1**

| | **Single-cell A1** | **Single-cell B1** |
|---|---|---|
| **Assembly statistics:** | | |
| Filtered CCS reads (bp) | 8,794,585,174 (44.2%) | 9,405,139,162 (46.6%) |
| Assembly size, primary (bp) | 598,293,718 | 454,096,399 |
| Assembly completeness | 19.4% | 14.7% |
| Contig N50 | 34,883 | 41,528 |
| Max contig size | 206,875 | 578,275 |
| Assembly size, alternative (bp) | 44,706,740 | 36,132,542 |
| Contig N50, alternative | 18,969 | 20,976 |
| Max contig size, alternative | 79,718 | 94,865 |
| **BUSCO gene models:** | | |
| complete | 1762 (12.8%) | 1236 (9.0%) |
| duplicated | 17 (0.1%) | 14 (0.1%) |
| fragmented | 58 (0.4%) | 250 (1.8%) |
| missing | 11960 (86.8%) | 12294 (89.2%) |
| **Mitochondrion:** | Complete | Complete |
